# Metagenomics-Toolkit: The Flexible and Efficient Cloud-Based Metagenomics Workflow featuring Machine Learning-Enabled Resource Allocation

**DOI:** 10.1101/2024.10.22.619569

**Authors:** Peter Belmann, Benedikt Osterholz, Nils Kleinbölting, Alfred Pühler, Andreas Schlüter, Alexander Sczyrba

**Affiliations:** IBG-5: Computational Metagenomics, Institute of Bio- and Geosciences (IBG), Research Center Jülich GmbH, Germany; Genome Research of Industrial Microorganisms, Center for Biotechnology (CeBiTec), Universitätsstrasse 27, 33615, Bielefeld, Germany; Computational Metagenomics Group, Center for Biotechnology (CeBiTec), Bielefeld University, Universitätsstraße 27, 33615, Bielefeld, Germany

**Author notes:** These authors contributed equally to this work (Peter Belmann, Benedikt Osterholz).

## Abstract

The metagenome analysis of complex environments with thousands of datasets, such as those available in the Sequence Read Archive, requires immense computational resources to complete the computational work within an acceptable time frame. Such large-scale analyses require that the underlying infrastructure is used efficiently. In addition, any analysis should be fully reproducible and the workflow must be publicly available to allow other researchers to understand the reasoning behind computed results. Here, we introduce the Metagenomics-Toolkit, a scalable, data agnostic workflow that automates the analysis of short and long metagenomic reads obtained from Illumina or Oxford Nanopore Technology devices, respectively. The Metagenomics-Toolkit offers not only standard features expected in a metagenome workflow, such as quality control, assembly, binning, and annotation, but also distinctive features, such as plasmid identification based on various tools, the recovery of unassembled microbial community members and the discovery of microbial interdependencies through a combination of dereplication, co-occurrence, and genome-scale metabolic modeling. Furthermore, the Metagenomics-Toolkit includes a machine learning-optimized assembly step that tailors the peak RAM value requested by a metagenome assembler to match actual requirements, thereby minimizing the dependency on dedicated high-memory hardware. While the Metagenomics-Toolkit can be executed on user workstations, it also offers several optimizations for an efficient cloud-based cluster execution. We compare the Metagenomics-Toolkit to five commonly used metagenomics workflows and demonstrate the capabilities of the Metagenomics-Toolkit by executing it on 757 metagenome datasets from sewage samples for an investigation of a possible sewage core microbiome. The Metagenomics-Toolkit is open source and available at https://github.com/metagenomics/metagenomics-tk.

## Introduction

Metagenomics addresses genome analyses of microbiome members residing in targeted environments and habitats. What complicates matters is that many microorganisms, especially in complex microbiomes, are currently unknown and often have not been cultivated yet. However, they may fulfill important functions in their respective ecosystem. These so far non-cultivable organisms are often referred to as the “microbial dark matter” [63]. Since the microbial dark matter represents a large fraction of microbiomes in almost all environments [43], large scale metagenomic analyses of thousands of samples have been carried out in environments like the ocean [55], soil [44] and human [59], to further explore these unknown organisms by generating metagenome assembled genomes (MAGs). It is to be expected that more large-scale studies will be conducted in the future due to the increasing amount of sequencing data [29].

Already metagenomic analysis of single samples generates a large amount of data and inherently needs substantial compute resources, especially for processing steps like metagenome assembly or annotation. While single samples can still be managed on single workstations, the processing of metagenomic data from complex environments, such as anaerobic digestion or sewage microbiomes, often involving hundreds or thousands of samples, requires a lot more resources. These can usually only be provided in the form of high performance or cloud computing services. Cloud computing has rapidly become an indispensable resource for researchers and organizations who have to store, manage, and analyze large amounts of data. The flexibility, scalability, and cost-effectiveness of cloud-based applications make it an ideal solution for large-scale metagenomics projects that require significant computing power and storage capacity. To use resources on cloud systems, appropriate computational workflows have to manage and scale on a plethora of virtual machines (VMs), so called cloud compute instances. The available resources should be used in an efficient way by specifying the required resources as close as possible to what is actually needed to reduce the costs in public clouds and facilitate the execution of multiple tools in parallel.

Available metagenomic workflows have their strengths and weaknesses and set their focus on a specific metagenomic analysis like the provision of multiple input types, or the optimization for specific computing environments. For example, MetaGEM [87] and MetaWrap [75] focus on a specific metagenomic topic, such as genome-scale metabolic modeling or a superior bin extraction algorithm. The MUFFIN [75, 76] workflow allows the incorporation of transcriptomic data and the nf-core/MAG [32] allows the combination of short and long reads and the incorporation of grouping information to perform co-assembly and binning. SequeezeMeta [74] is a software that can run on small desktop computers. However, these workflows are either not designed for cross-dataset analyses, such as dereplication on thousands of samples, or they are not optimized for cloud-based cluster systems, which can limit their scalability.

Apart from the need for a new efficient cloud-enabled solution with additional analytical capabilities, we set a focus on the ability to re-analyse publicly available datasets with updated tools. The Critical Assessments of Metagenome Interpretation CAMI 1 and 2 initiatives [70, 47] have highlighted advances in different metagenomic analysis strategies. A re-analysis using state-of-the-art tools should provide improved insights into microbiomes.

The analysis or re-analysis of metagenomic samples, especially on a large-scale, sets an emphasis on the explorability of the data. Analysis results from hundreds or thousands of samples should be easy to explore and comprehend, especially for users without a background in computer science. For this user group, computed results should not be solely available in the form of text files. The explorability becomes even more important in the case of comparative analyses. Comparative analyses of predicted coding sequences, their annotation, biological processes or abundances of MAGs will become slow and tedious without a sufficiently fast database engine and suitable visualization.

To enable enhanced reproducibility and scaling capabilities, we developed a Nextflow [17] workflow which tackles the aforementioned challenges, allows the application on single workstations and optimizes the application in cloud environments and enables automatic (re-)analysis of public datasets. We refer to this workflow as the “Metagenomics-Toolkit” or “Toolkit” for short. The Metagenomics-Toolkit offers novel analysis capabilities compared to other workflows in the form of sample-wise consensus-based plasmid detection and fragment recruitment, as well as cross-dataset dereplication and co-occurrence analysis enhanced by metabolic modeling. To broaden accessibility and usability for users without a computer science background, the Toolkit is also available through a user-friendly web-based interface, powered by the Cloud-based Workflow Manager (CloWM) [18] service. Toolkit outputs can be investigated via the Exploratory MetaGenome Browser (EMGB) [22] web application to collate, integrate and visualize all Toolkit results in a user-friendly, graphical format.

In addition, we applied a machine learning approach to predict RAM requirements of an assembler based on the characteristics of the input dataset. This allows more precise resource distribution, which may result in a reduction of the requested RAM and, in certain instances, the elimination of the necessity for dedicated high-memory hardware. Furthermore, this method could be adapted to other bioinformatics tools in the future to optimize their resource consumption.

To demonstrate the various analysis capabilities of the Toolkit, we reanalyzed metagenomic datasets of untreated sewage, mainly collected by the Global Sewage Surveillance project. The examination of metagenome datasets from untreated sewage samples allows, for example, to monitor the distribution of genes of interest, such as antimicrobial resistance genes (AMR) [50]. Here we focus on the detection of the sewage core microbiome, i.e. species with a global distribution, and most importantly, the full reproducibility and automation of our analysis, which will be useful for continuous tasks such as the monitoring of AMR or pathogenic organisms on a global scale. All results generated by our tools are publicly available for further investigation (see Data Availability).

## Materials and Methods

The Metagenomics-Toolkit is built upon the Nextflow workflow engine, designed to streamline and automate the large-scale processing of metagenomic datasets. For the purposes of our analysis, we deployed Nextflow on a cluster within the de.NBI Cloud [7] infrastructure using BiBiGrid [1], an open-source cluster management tool. This setup allowed us to efficiently handle the complex computational demands of the sewage core microbiome analysis. In this chapter, we provide a detailed account of the configuration and tools used for this analysis, as well as the machine learning strategy we implemented to predict the RAM requirements of the assembler used.

### Distributing bioinformatics workloads with Nextflow

Processing large amounts of metagenomic data manually can be complex, time-consuming and is prone to errors. Consequently, it is unreasonable to process multiple metagenomes simultaneously without some sort of automated connection and execution of individual computational processing steps, as e.g. provided by workflow engines.

The Toolkit is written in the domain specific language (DSL) Nextflow, which was developed to support the writing of data-intensive computational workflows especially in bioinformatics. Nextflow follows the dataflow programming model, which implicitly defines parallelisation through in-and output declarations. The resulting applications are therefore inherently parallel. This makes it possible to distribute tasks across multiple VMs. Nextflow also supports containerization technology, such as Docker, which makes it possible to share and run workflows on different systems, so called executors, like workstations, cloud-based services, and high-performance computing clusters without the need for manual dependency management on different computing environments. Many executors, like SLURM [83], Kubernetes and AWS Batch, are supported by Nextflow. Workflows launched on said executors are able to read their inputs and write their results directly to object storage systems that are compatible with the Amazon S3 API and their submitted commands are automatically resubmitted in the event of errors.

For the Toolkit, the newer DSL2 was used, which enables the subdivision in a more modularized workflow structure. Tools and sub-workflows of major workflow steps are separated in their own module files.

### Cluster management in cloud infrastructures using BiBiGrid

In order to perform our large-scale sewage microbiome analysis via the Metagenomics-Toolkit using the de.NBI Cloud infrastructure, we utilized BiBiGrid, an open-source tool for setting up and managing clusters in cloud environments.

For our use case, BiBiGrid will set up a cluster for grid computing, utilizing the SLURM workload manager. Figure 1 shows our simplified BiBiGrid setup, with a “master” VM used to submit jobs and execute the Nextflow binary and two “worker” VMs tasked with executing the submitted workflow commands. The SLURM workload manager distributes self-contained units of work, called jobs, across a cluster, allowing one or multiple jobs to run simultaneously on a single worker. Across all worker VMs, BiBiGrid will also set up a shared file system based on NFS for Nextflow’s working directory, that collects intermediate results. In addition, a “scratch” disk is available on each worker, which is located on a hard drive on the host of the respective worker VM.

**Fig 1.**
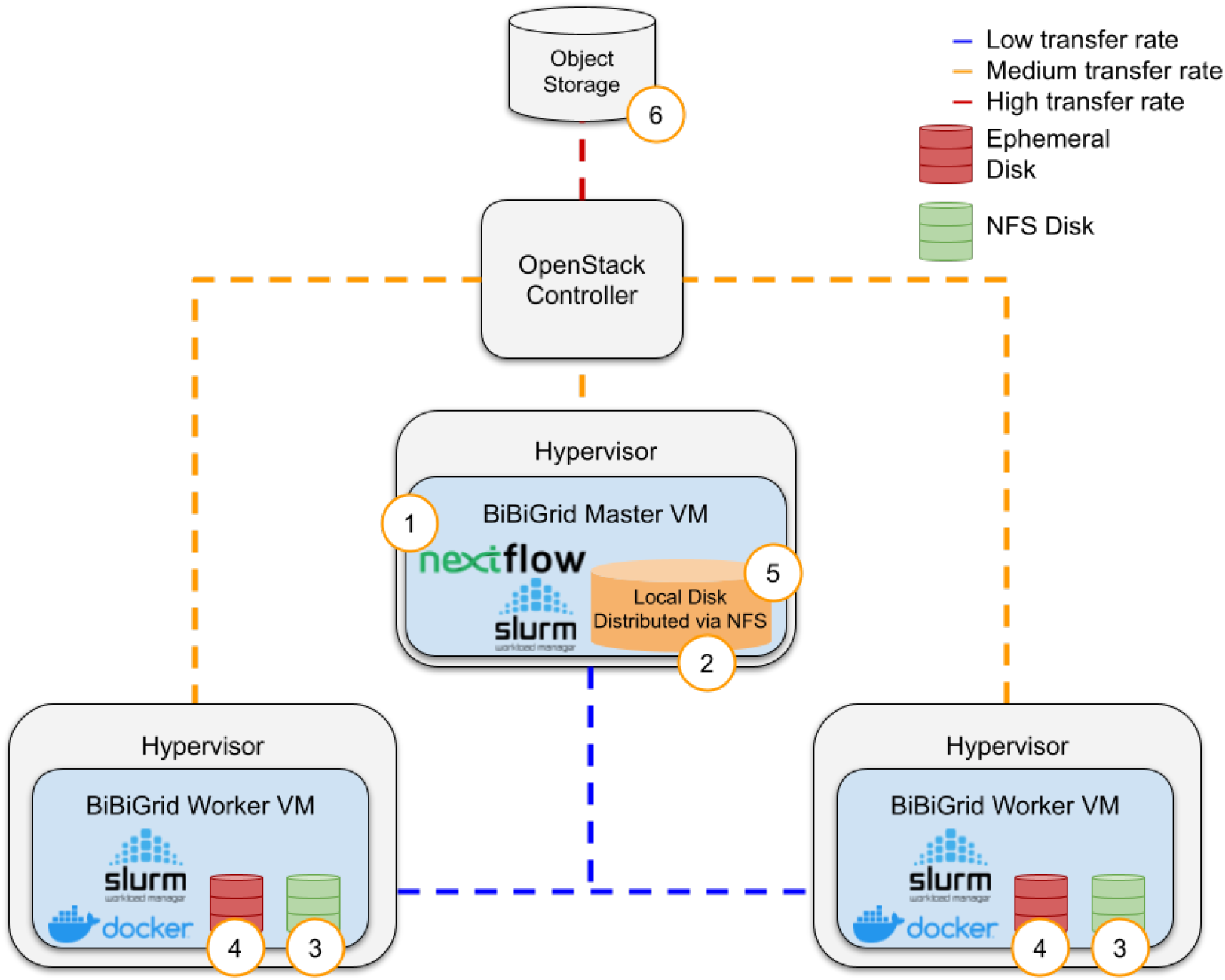
The figure illustrates the data handling of the Metagenomics-Toolkit deployed on a simplified BiBiGrid cluster, comprising one master virtual machine (VM) and two worker VMs, modeled after the characteristics of a de.NBI Cloud site. The numbers represent typical steps in the execution of the Metagenomics-Toolkit. 1) The Master VM is used to start a Nextflow workflow via SLURM. 2,3) The Worker VM is used to store intermediate workflow results on a NFS. 4) An ephemeral disk is used to store intermediate results of a single command. 5, 6) Results of individual commands are stored on the NFS, while final results are transferred to the object storage.

Another feature of BiBiGrid is the ability to auto-scale the cluster. The used SLURM Workload Manager is able to monitor the current workload and, provided there are enough resources, spawn new instances as needed to keep up with demand. Additionally, when the workload decreases, SLURM can automatically terminate instances to reduce costs.

### Technical characteristics of the de.NBI Cloud infrastructure

For our analyses, we leveraged the de.NBI Cloud infrastructure, which consists of multiple independent cloud sites, each powered by OpenStack, an Infrastructure as a Service (IaaS) platform. OpenStack allows for the management of key cloud computing components such as storage, networking, and compute resources. In addition, the de.NBI Cloud infrastructure offers SimpleVM, an abstraction layer on top of OpenStack. Using either OpenStack or SimpleVM we configured virtual machines (VMs) with customizable settings, known as flavors, to define parameters such as CPU count and RAM capacity. To present further characteristics of a de.NBI Cloud site we take a cluster setup consisting of one master VM and two worker VMs as an example, as illustrated in Figure 1. The master VM orchestrated the workflows, while the SLURM workload manager allocated individual tasks and their associated data to the available worker VMs. However, a limitation of this setup is the shared bandwidth among VMs, which can cause high latency when multiple processes, potentially running on hundreds of VMs, concurrently access a shared file system. This synchronization of data between the master and worker VMs may result in reduced data transfer rates. In comparison, data transfer between a VM host, called Hypervisor, and OpenStack’s object storage, which is accessed through controller nodes, provides better connectivity and higher transfer rates. Object storage is used to store input and final output data. To mitigate some of these performance bottlenecks, Hypervisors are equipped with local SSD drives as ephemeral disks providing the fastest data read and write speeds.

### Preparation, feature selection and machine learning model assessment for the peak RAM prediction

To optimize the parallelization of the Toolkit, we applied a machine learning approach to estimate the peak RAM consumption of the MEGAHIT assembly. The resulting product of this approach is a model, i.e. a mathematical representation of the knowledge learned from the data, where the process of feeding data to a machine learning algorithm is called training. In the following subsections we explain the data preparation and feature selection for training and testing a machine learning model. In a final subsection we describe the assessment of the model.

#### Data preparation for training and testing a machine learning model

MEGAHIT applies a multiple k-mer size strategy, where multiple assemblies based on different k-mer sizes are constructed. Our hypothesis is that diversity and sequencing depth of the biological sample are the main factors that influence MEGAHIT’s memory consumption. To predict the peak memory, 1212 metagenome datasets from environments of varying complexity, i.e. soil, biogas reactors and nasopharyngeal, were assembled twice, using MEGAHIT’s default and meta-sensitive parameter settings. In the latter case, a wider range of k-mer sizes as compared to the default setting is used, leading to a more accurate and complete assembly, but also to a higher RAM consumption. The peak total amount of memory used by MEGAHIT was monitored by Nextflow and extracted from Nextflow’s trace file.

We applied 10-fold cross-validation for every model, where the training set itself was randomly split in multiple folds consisting of a validation and test subset. Finally, we compared different regression models, then fine-tuned and evaluated the best performing model.

#### Selecting suitable features for the prediction of peak memory

We started by inspecting the following features: Number of bases, GC content, minimum read length, average read length, total number of reads, and the k-mer counts of size 13, 21 and 71 of all datasets produced by K-mer Counter (KMC). Based on the resulting k-mer counts, we extracted the following parameters: Total number of k-mers, total number of distinct k-mers, mean, standard deviation, maximum of the total number of k-mers per k-mer frequency, and the sum of the lowest 5% (*Quantiles 5 k-mers*) and highest 5% (*Quantiles 95 k-mers*) of the total number of k-mers per k-mer frequency. In addition, we included the Nonpareil community diversity index, which summarizes the redundancy or uniqueness of sequences within a given dataset.

To reduce the number of features needed to predict actual peak RAM usage, we examined the Pearson correlation coefficient between the variables and the peak RAM consumption (Supplementary Figure 1). For generating different machine learning models, we selected 18 features with a Pearson correlation coefficient greater than 0.6 (p-value *<* 0.05), as follows: GC content, Nonpareil diversity index, total number of reads, minimum and average read length, total number of bases and k-mer counts of size 13, 21 and 71. Based on k-mer counts we extracted the following features: Total number of k-mers (k-mers 13, 21, 71), total number of distinct k-mers (k-mers 21, 71), mean (k-mers 21, 71), standard deviation (k-mers 21, 71), maximum of total number of k-mers per k-mer frequency (k-mer 13), sum of the lowest 5% (*Quantiles 5 k-mers*, k-mers 13) and highest 5% (*Quantiles 95 k-mers*, k-mers 13, 21, 71) of the total number of k-mers per k-mer frequency.

#### Assessing machine learning models and reducing the final set of features

We applied several regression-based machine learning methods, namely linear regression, support vector machines, decision trees, random forest and extremely randomized trees. Based on the mean and standard deviation of each root mean square error (RMSE) of the cross-validated sets, we compared the resulting models (Supplementary Table 3). Based on these comparisons, we chose the extremely randomized trees regressor, which had the lowest mean and standard deviation of RMSE in the default model and the third lowest mean in the meta-sensitive model, respectively, and the lowest standard deviation among the best three models. In addition, the extremely randomized trees regressor allows to easily retrieve feature importances and thereby conduct feature selection. As a next step, we optimized the hyperparameters, using an exhaustive grid-search over provided parameters, like the minimum number of samples at a leaf node.

### Available tools and configuration details for the sewage core microbiome analysis

The Toolkit offers several tool options within each category (assembly, binning, etc.). For convenience, we provide a list of all available tools in Table 1 and provide a more detailed table in Supplementary Table 4, that also highlights tools used specifically for the analysis of the sewage core microbiome analysis. Regarding the core microbiome calculation, we will focus on the mapping configuration part, which is important for the selection of the core microbiome members while the final configuration file can be found in the “Data Availability” section. We took all SRA run accession IDs of all sewage samples from a previous publication [49] and extended them with a region assignment according to World Bank country groupings (Supplementary Figure 5), similar to the groupings made by Jespersen et al. 2022 [25].

**Table 1.** List of all tools that are available in the Metagenomics-Toolkit. Tools that are used in multiple modules have the module name “Multiple Modules”.

| Modules | Tools |
| --- | --- |
| Annotation | MMseqs [20], MMSeqs taxonomy [48], GTDB-tk [10], CheckM [58], Prokka [71], Prodigal [24], RGI [2] |
| Assembly | Flye [31], Assembler Resource Estimator, Spades [52], MEGAHIT [37] |
| Binning | MetaBAT2 [28], MetaCoAG [46], Metabinner [80], MAGScoT [67] |
| Co-Occurrence | Spiec-Easi [33], igraph [14], further R libraries |
| Dereplication | Pyani [62], SANS [82] |
| Genome-scale metabolic modeling | CarveMe [45], MEMOTE [41], SMETANA [84], GapSeq [86] |
| Input | Pysradb [12] |
| Multiple Modules | Minimap2 [39], Bowtie2 [35], Mash Screen [53], [77], Seqkit [72], SAMtools [15], CoverM [5], BWA-MEM [38], BWA-MEM2 [78] |
| Plasmids | Platon [69], ViralVerify [4], MOB-suite [64], SCAPP [61], PlasClass [60], PLSDB [68] |
| Quality Control | Fastp [11], Porechop [81], Filtlong [21], Nanoplot [16], KMC [30], Nonpareil [65] |

We removed all negative controls from the 951 samples and selected only one sample from each biological replicate. This resulted in 757 sewage samples with an average size of 12.9 Gbp before and 11.6 Gbp after quality control by running the Toolkit on two de.NBI Cloud sites. After assembly, binning and dereplication, we mapped each sample against a dereplicated set of 3473 MAGs that are at least 50% complete and are at most 5% contaminated according to CheckM. Completeness and contamination are calculated by counting lineage-specific marker genes. We require that at least 90% of the genome is covered by 1-fold base coverage. Supplementary alignments are excluded and mapped reads are only accepted if the read is at least 95% aligned with 95% identity. The final abundance table can be inspected in Supplementary Table 5.

## Results

### Metagenomics-Toolkit: An overview of its design and implementation

The Metagenomics-Toolkit combines several well established bioinformatics tools and methods by using the workflow system Nextflow to create related modules and combining them in novel ways. A short list of all available tools can be found in Table 1, a more detailed version in Supplementary Table 4 and a summary of all modules and their connections can be found in Figures 2 and 3. Before detailed explanations are given in subsequent chapters, this chapter provides an overview of the Toolkit modules and their functionality.

**Fig 2.**
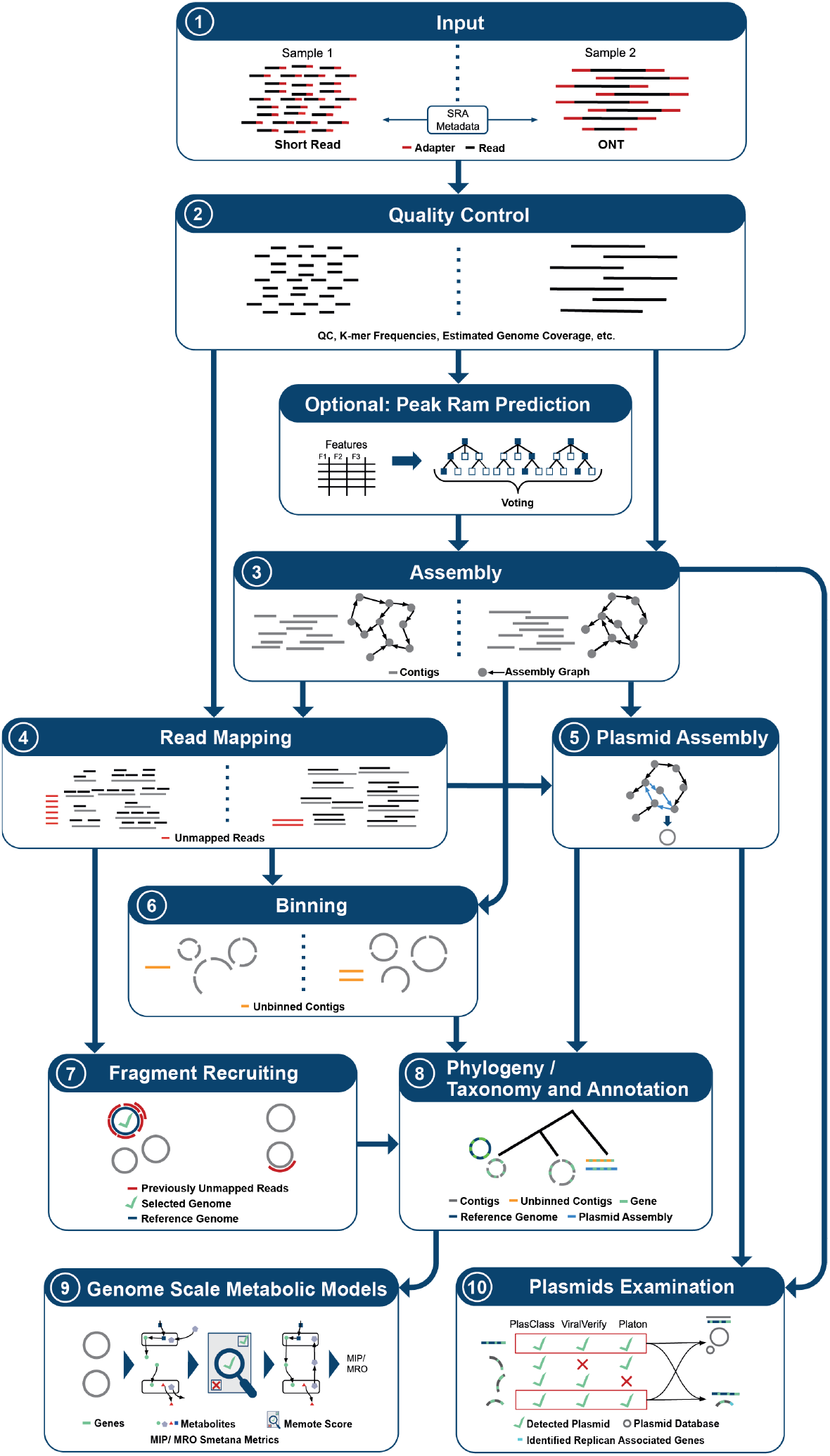
A simplified overview of the Metagenomics-Toolkit single sample workflow. The processing of different kinds of reads in the main modules of the Toolkit is illustrated in a step-by-step manner, from top to bottom. The split modules illustrate the different processing methods for short reads on the left and long reads on the right of a module. The metadata of the input reads is checked against the SRA metadata to determine whether ONT or Illumina is provided when SRA is used as a resource for input files (1). All reads are then first quality controlled (2). If Illumina reads are provided and MEGAHIT is selected as the assembly tool, then the peak RAM is predicted. After assembly (3), the contigs are provided as input to the read mapping module (4) and the assembly graph is used for plasmid assembly (5) and optional binning (6). Reads that could not be mapped back to the assembly are mapped against a set of genomes provided by the user (7). All contigs, including the plasmid assembly and genomes detected in the fragment recruitment step, are used as input to the annotation module (7). All contigs, also those from the plasmid assembly, are provided to the plasmid module (10) for further analysis. Predicted proteins are used as input to the genome-scale metabolic modeling module (9).

**Fig 3.**
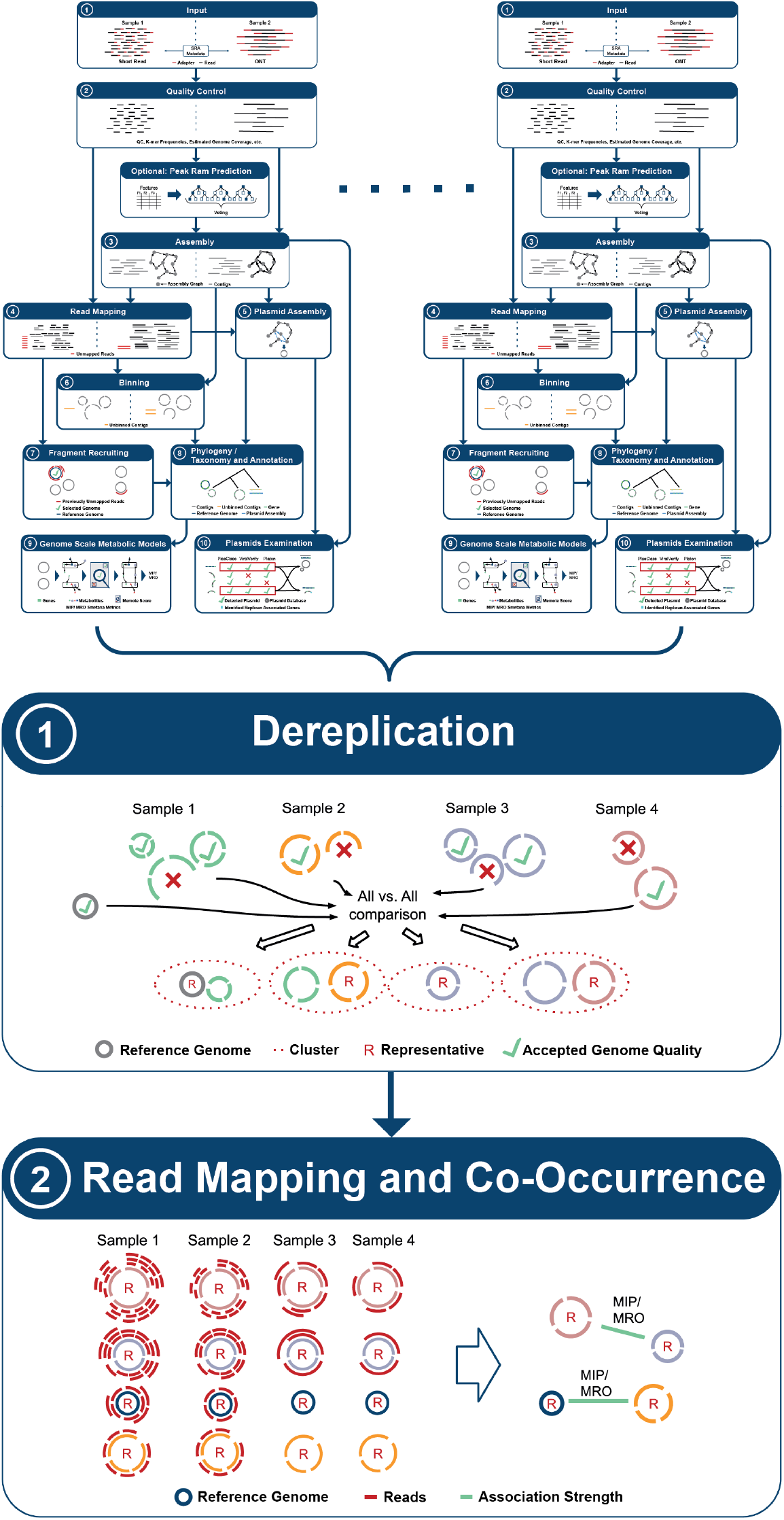
A simplified overview of the Metagenomics-Toolkit aggregation part. Once the single sample workflow has been completed, the aggregation of all datasets is initiated. As illustrated in the figure, the results of single sample workflows serve as input for the aggregation workflow. Redundant species MAGs are dereplicated (1) in order to obtain a unique set of representative species MAGs. The reads of all samples are mapped against the representative MAGs (2). Based on the abundance values, the co-occurrence of the representative MAGs is calculated. The edges of the resulting co-occurrence network are annotated with the MIP and MRO metrics.

In addition to the functionality of running all parts of the workflow consecutively, the Toolkit is subdivided into so-called modules that can be executed separately. In most cases, modules contain multiple tools for the same or similar type of application. The user can select the appropriate tool for a particular dataset based on personal preferences or the findings of benchmarking projects such as the CAMI challenges [70, 47]. The workflow accepts a path, link or S3 address to files containing paired-end or Oxford Nanopore sequences as input.

In general, the Toolkit follows a two-step strategy for generating MAGs. In a first step, hereafter referred to as the “per-sample” step, all samples are assembled and binned independently, which is expected to result in higher quality genomes compared to a co-assembly approach [70]. Subsequently, in a second step, called aggregation, all MAGs are dereplicated into clusters at the species or strain level [59]. MAGs that have been dereplicated in the second step are analyzed for possible associations using a co-occurrence approach. Both steps can be performed in one or separate calls. The advantage of this step-wise approach is the option to process a large number of samples independently, allowing the use of multiple independent compute infrastructures. The optional aggregation can be done by providing the output of the per-sample step as an input to the second one.

Alongside the technical features outlined in the following chapters and the standard functionality commonly found in metagenomics workflows, such as assembly, binning, and annotation, the Metagenomics-Toolkit offers distinctive functionalities. These functionalities serve two particular purposes: first, the automated re-analysis of publicly available datasets, and second, the enhancement of the analysis of MAGs, plasmids, and metagenomic datasets as a whole, which will be the focus of the following sections.

#### Reconstruction and annotation of MAGs

The Toolkit can either be executed in its short or long read mode, either automatically determined by evaluating corresponding metadata or manually specified by the user. The choice of the mode affects multiple parts of the workflow as illustrated in Figure 2 (Points 2 -7) and Figure 3 (Point 2).

In its short read mode, raw short reads are subjected to quality control (Figure 2, Point 2) by using Fastp for adapter removal and quality trimming. Additionally, k-mer count frequencies are generated using KMC and the Nonpareil diversity using the tool ‘Nonpareil’ is determined. Both values are necessary for subsequent analysis steps if peak RAM prediction mode is enabled (Figure 2, optional). In long read mode (ONT) [79], Porechop is used for the removal of adapter sequences and Filtlong for read trimming.

While in short read mode, the user can choose to assemble reads using metaSPAdes or MEGAHIT (Figure 2, Point 3), in long read mode metaFlye is offered. To address variable error rates in Oxford Nanopore sequencing, metaFlye parameters for specifying the expected error rate are automatically determined based on the median PHRED quality score. Resulting assembly graphs are passed for their usage to other modules like plasmid assembly via SCAPP (Figure 2, Point 5) or binning in MetaCoAG (Figure 2, Point 6).

Preprocessed reads are mapped back to the generated contigs using Bowtie2 or BWA-MEM2 for short reads and Minimap2 for long reads (Figure 2, Point 4). The obtained read coverage information of these mappings is utilized for the next binning step and for the generation of assembly coverage statistics using CoverM. MetaBAT2 is the default binner for both strategies, but can also be exchanged with Metabinner in short read mode or MetaCoAG in long read mode (Figure 2, Point 6).

Prokka annotates all MAGs, all contigs that could not be binned, and the plasmids that were assembled separately (Figure 2, Point 8). Prokka utilizes a set of predefined Toolkit parameter settings as well as parameter settings that depend on the taxonomic classification of the input sequence. Here, the taxonomy kingdom of the MAG classification using the Genome Taxonomy Database Toolkit (GTDB-Tk) is passed to Prokka for the selection of the correct annotation mode. MMSeqs taxonomy is used to assign taxonomic labels to all predicted genes (Figure 2, Point 8).

Predicted coding sequences are further annotated by the Resistance Gene Identifier (RGI) and MMSeqs (Figure 2, Point 8). RGI predicts antibiotic resistance genes by conducting a homology search against the Comprehensive Antibiotic Resistance Database (CARD) [2]. As stated in a recent article by Papp, M. and Solymosi, N., CARD focuses on acquired resistance genes and antimicrobial resistance-associated mutations, and therefore is, for a variety of study settings, the preferable choice in comparison to other antimicrobial resistance gene databases [56].

MMSeqs [48, 73] is used to search for homologous protein sequences in databases such as KEGG [27] for functional annotation, BacMet [54] for antibacterial biocide or metal resistance and VFDB [42] for virulence factors. If sufficient memory is available, all databases are stored in RAM to accelerate computations. It is possible to add additional databases according to user requirements. One example is MetaCyc [9], an alternative metabolic pathway database. The user can provide the database as an HTTP/S, S3 link or a local file path, as long as the database is compressed according to the Zstandard [13]. To accelerate the annotation module execution, a divide and conquer pattern is applied. The annotation is performed separately on MAGs, unbinned contigs, and assembled plasmids, in parallel for every database.

#### Identification of unassembled community members

Current state-of-the-art metagenomic tools are limited in their ability to assemble and bin the entire microbial community, i.e. potentially important genomes may be missed [70, 47]. To address this issue, a fragment recruitment strategy is used to detect known genomes that are part of the community but could not be assembled or binned (Figure 2, Points 4, 6 and 7). First, all reads failed to map to the assembly are screened against a user-provided database of reference genomes. As this can get computationally time-consuming, if, for example, all representative genomes of the GTDB Taxonomy [57] are used, Mash Screen is run as a fast preliminary filter step. This reduces the possible search space by limiting the number of genomes that have to be checked in the next, more computationally extensive step. Here, identified datasets containing the detected genomes are aligned against the Mash Screen matches using BWA-MEM2 for short reads and Minimap2 for long reads. Finally, alignments are inspected using CoverM. Genomes are reported as a final match if they meet a user-defined percentage threshold of base coverage (default: 90%). All this way identified reference genomes are then used as additional inputs for all proceeding modules, such as dereplication.

#### Consensus-based Plasmid identification

The detection of plasmids within metagenomic datasets can be achieved, in general, by two methods. One approach involves identifying plasmid-specific genes and proteins which can be done via the tools Platon, ViralVerify and PlasClass. These tools are executed on all contigs of the preceding assembly process. Only contigs for which all specified tools agree are reported as possibly belonging to plasmids to increase the precision of plasmid detection (Figure 2, Point 10). Another way to perform plasmid detection is by assembling them using SCAPP, that builds upon the assembly graphs of MEGAHIT, metaSPAdes or metaFlye. Finally, detected plasmids are further analyzed. Possible similar plasmids are searched in the PLSDB, to distinguish between previously reported and novel plasmids. Other characteristics, like the predicted host range of the plasmid, are analyzed using MobTyper.

#### Co-occurrence analyses and metabolic modeling

The co-occurrence module, which is part of the aggregation step (Figure 3, Point 2), allows users to analyze co-occurring organisms in different datasets based on their per-sample occurrence and abundance. By applying, for example, network theory as part of the downstream analysis on the resulting co-occurrence networks, valuable insights into the complex structure of microbial communities can be gained. In human-related microbiome datasets, co-occurrence allows to infer the influence of co-occurring organisms on the host’s health [36]. Due to the inherent difficulty in interpreting the resulting co-occurrence networks [66], metrics derived from genome-scale metabolic modeling (Figure 2, Point 9) are included in the final co-occurrence network. In the following section, we will first describe the functionality of the independent modules and finally explain the integration of genome-scale metabolic modeling and co-occurrence.

MAGs of different samples that were generated in the per-sample step of the workflow are assigned to species or strain clusters by applying a hierarchical strategy adopted from Pasolli et al. [59] (Figure 3, Point 1). First, all generated MAGs are filtered by completeness and contamination and then pre-clustered via Mash. As in the fragment recruitment step, Mash distances of all MAGs were used for an average linkage clustering due to its fast distance calculation. Clusters were formed by using a 95% cutoff. In a second step, a representative genome is selected based on a scoring system that considers CheckM completeness, CheckM contamination, CheckM heterogeneity, N50 and coverage for each cluster. Additionally, a slower but more accurate Average Nucleotide Identity (ANI) computation is conducted between all representatives in order to improve cluster formation. In case the ANI of two representatives is above 95%, their respective clusters are merged.

Quality-controlled reads from all samples are mapped against all representative genomes to determine the abundance of each genome in each sample (Figure 3, Point 2). Based on the compiled abundance table, possible associations between MAGs (Figure 3, Point 2) can be calculated using two different approaches. The first approach uses pairwise Spearman’s non-parametric rank correlations. Specifically, p-values are calculated for correlations between each pair of MAGs for multiple permutations of the abundance table. Adjusted p-values are then obtained using the Benjamini-Hochberg procedure [8]. Finally, only reliable associations are used based on the adjusted p-values. The second approach is based on co-occurrence networks computed from genome abundances extracted from 16S ribosomal RNA gene datasets. Here, Spiec-Easi is applied on the abundance table to infer an underlying graphical model using the concept of conditional independence. The resulting associations between MAGs, regardless of the chosen approach, represent a co-occurrence graph whose nodes are further annotated by using the GTDB taxonomy. The graph can then be analyzed by using network theory and plotted by Python/R libraries and tools like igraph or gephi [6].

The Genome-Scale Metabolic Modeling module generates genome-scale metabolic models (GEMs) that are mathematical representations of chemical reactions within a microbial organism [3]. GEMs are generated from the corresponding annotation of all high quality MAGs via CarveMe or GapSeq and quality controlled using MEMOTE (Figure 2, Point 9). Further, possible cases of cross feeding and competition between microbiome members per sample are assessed by running SMETANA. SMETANA outputs metrics like the metabolic interaction potential (MIP) and the metabolic resource overlap (MRO). Depending on the available amount of computational resources, the user can enable the computation of the species coupling score (SCS), which measures the dependency of the growth of a given species in the presence of another species in a community. MRO, MIP and SCS allow the researcher to make assumptions about the degree of a possible interaction between microbial community members.

To facilitate the interpretation of co-occurrence networks, the MRO and MIP metrics are computed for each pair of MAGs that are connected by an edge in the co-occurrence network. A co-occurring pair of MAGs with a high MIP might be a hint for cross feeding, while a high MRO value might suggest organisms competing for the same resources.

### Optimizing efficiency for cloud-based cluster system

While it is possible to run the Toolkit on a single user workstation, the key design decision for the Toolkit was to use cloud-based compute clusters that allow workflows to scale in cloud environments using proven job workload managers like SLURM. Figure 1 illustrates an exemplary setup that we have utilized using the BiBiGrid open-source tool for a cluster configuration inside a cloud environment. The following sections describe the optimizations that have been implemented to make the Toolkit particularly compatible with cloud-based compute environments in general and specifically with the setup of the de.NBI Cloud, whose characteristics and challenges are described in detail in the “Materials and Methods” section and visualized in Figure 1. In essence, due to bandwidth limitations, software should reduce the data transfer between VMs and instead use the local disk and object storage for input, output and intermediate results.

#### Efficient and automated input data and database download

The Toolkit enables automated and streamlined processing of publicly available datasets by specifying a list of SRA or study IDs that are automatically retrieved either directly from NCBI or from a user-defined mirror. Individual datasets can be processed by providing a list of sample names and paths to their respective locations, either locally, by HTTP/S or S3. Prior to any dataset download, correctness and accessibility of SRA paths are verified, along with any associated metadata for public datasets, which is automatically downloaded along.

In the default execution of Nextflow runs, all input files are stored on the Network File System (NFS) with all intermediate results (Figure 1, Points 2 and 3). This can lead to bottlenecks if too many read/write operations occur at the same time. This can become even more problematic when large sequence databases, such as NCBI-nr, are downloaded to the NFS and queries to these centrally provided databases are distributed across multiple compute instances. To reduce the load on the NFS, the workflow has been optimized to handle file downloads more efficiently. In contrast to the default execution of Nextflow runs, the input fasta files are downloaded directly to the scratch disk of the cluster instance that requires them (Figure 1, Point 4). This also reduces the need for large amounts of space on the NFS. Keeping local copies of reference databases on the scratch disk on each instance is more efficient in the de.NBI Cloud setup. To ensure consistency between all local copies, the Toolkit integrates an automated task for this purpose. Databases can be uploaded to an S3 compatible object storage and referenced in the configuration of every tool, respectively, in the form of a S3 path and database-specific md5sum hash. During the execution of a tool, required databases are searched in a predefined directory that is located on the scratch disk, and its md5sum hashes are checked. If the database is not found or the hashes do not match, i.e. the database is missing or present in a different version, it will be automatically downloaded from the S3 cloud storage and stored on the scratch disk. This ensures that the configured version of the database is always available for analysis. The Unix flock locking process is used to ensure that multiple downloads of the same database are not started at the same time on the same worker node, since each database download is a separate job in the context of SLURM (see Material and Methods). The Toolkit can access a large number of databases that are relevant for metagenomic analyses and have been uploaded to the de.NBI Cloud Bielefeld object storage. Corresponding links and checksums are provided in the default Toolkit configuration file (see Data Availability).

#### Tool execution on cloud instances

For tool execution, the Toolkit utilizes containerization by using Docker containers and thus allows for the convenient distribution and execution of software along the worker instances of a cluster without the need for manual installation. When available, we used public Docker images created by the Bioconda community [19], otherwise we created our own (Supplementary Table 4). To avoid high input/output (I/O) activity on the shared NFS which can lead to a high latency, each tool is executed in the scratch directories of the cloud instances. Only results are copied from the scratch directories to the shared NFS. An additional mode allows further reduction in the NFS usage, by circumventing the normal Nextflow SRA input download mechanism, with all remote files being placed directly into the worker scratch directory instead of the usual Nextflow working directory. By using this mode, the raw files do not consume additional disk space on the shared file system and the download can be performed in parallel on the worker nodes.

#### Provenance and Visualization

In order to ensure the provenance of our workflow, i.e. the structured storage of the workflow execution details, the workflow stores each command, as well as results, in directories named in accordance with the module name, module version, executed tool and its version number (Supplementary Figure 4). In conjunction with Docker containerization, this approach warrants reproducibility.

**Fig 4.**
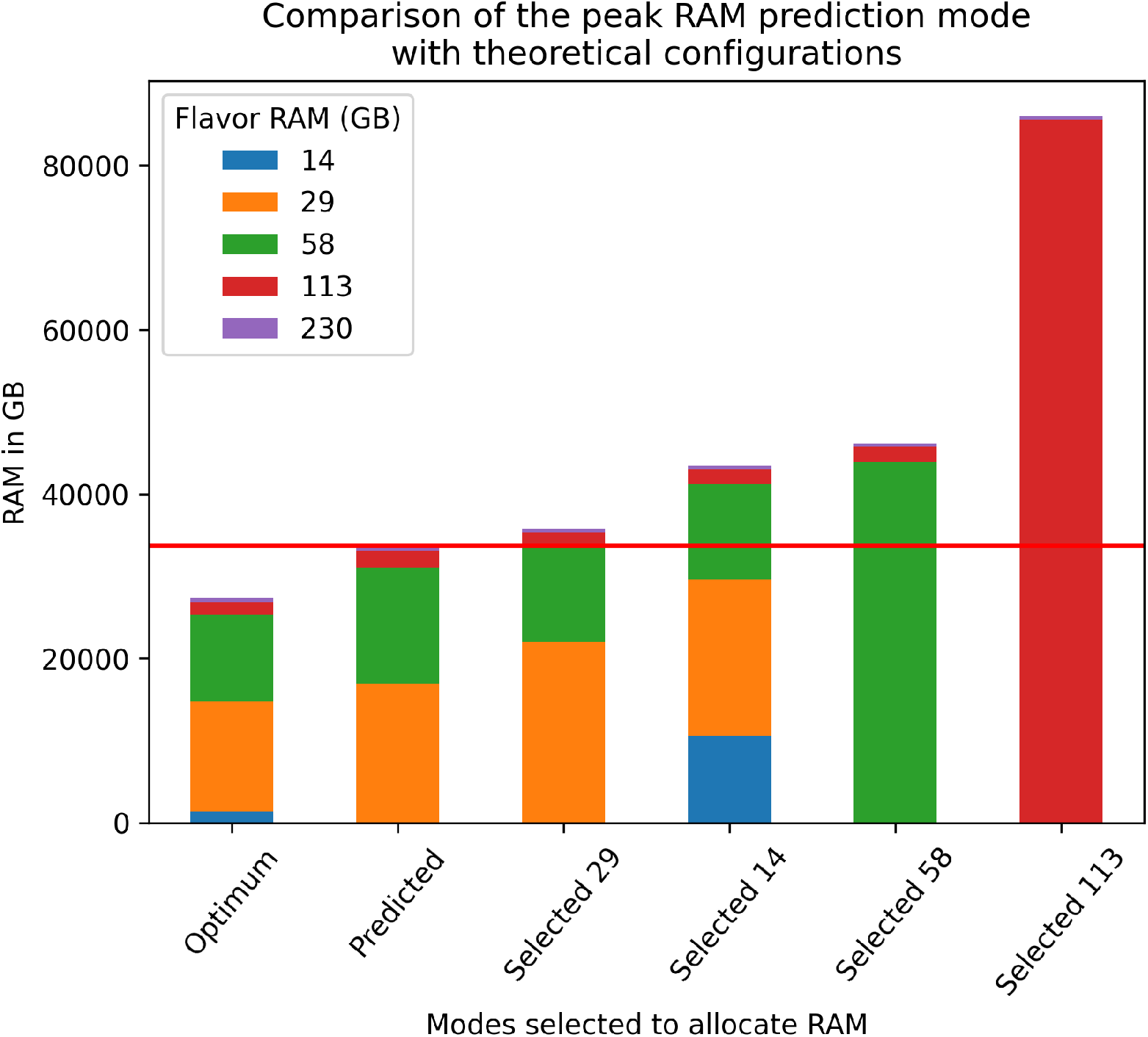
Different RAM configurations when assembling 757 sewage samples. Selected* configurations are theoretical settings, where a user has selected a flavor for all datasets and the workflow only increases the flavor if the assembler fails due to insufficient RAM. The Optimum mode uses the most appropriate flavor for every dataset. The Predicted mode uses the predicted flavor for the respective sample. All modes are calculated using the peak RAM value reported by Nextflow and the predicted peak RAM value for all samples. The total peak RAM value of the predicted mode is represented by a red line.

Standardizing the output of multiple modules simplifies the use of the Toolkit as part of other workflows and allows the output of the per-sample step to be reused as input for the aggregation step. Standardization in both cases is particularly useful when tools of the same module are being replaced.

The Toolkit output can be transformed into an EMGB compatible input by running a post-processing script on the Toolkit output files, to create json files containing assembly, binning and annotation information. EMGB is a web interface that can be used for the visual exploration of metagenomic datasets. Large datasets containing millions of genes and their annotations are pre-processed and visualized, to be searched in real-time by the user. The platform provides access to different aspects of one or more datasets via an interactive taxonomic tree and dynamic KEGG metabolic maps for each dataset, allowing researchers to explore their datasets at the level of genes, contigs, MAGs, pathways or biological process statistics (Supplementary Figure 6).

**Fig 5.**
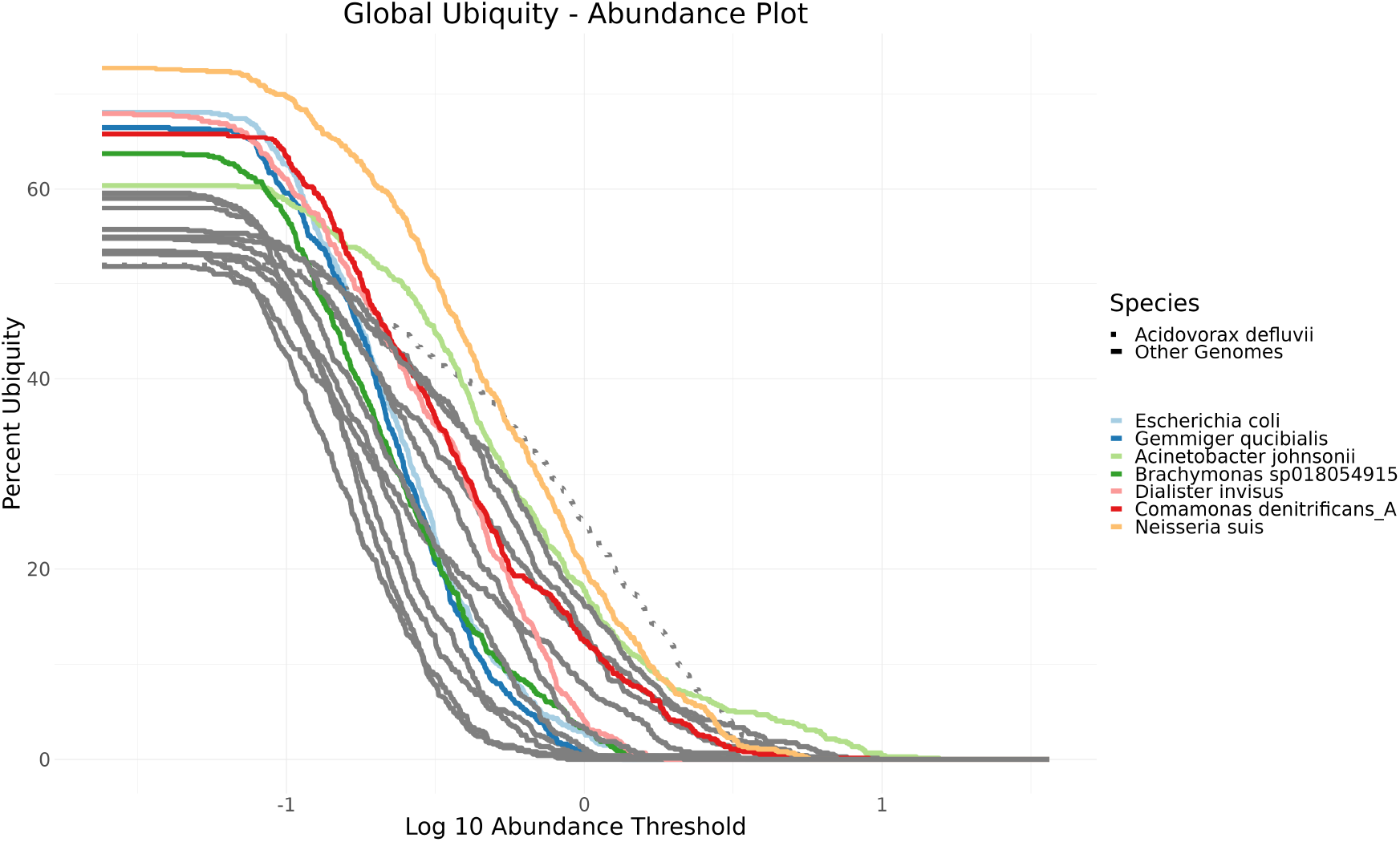
The Ubiquity - Abundance plot shows for each species its occurrence in samples (Percent Ubiquity y-axis) with a specific minimum abundance (Log10 Abundance Threshold x-axis). Only species with at least 50% ubiquity are displayed and species with more than 60% are colored.

**Fig 6.**
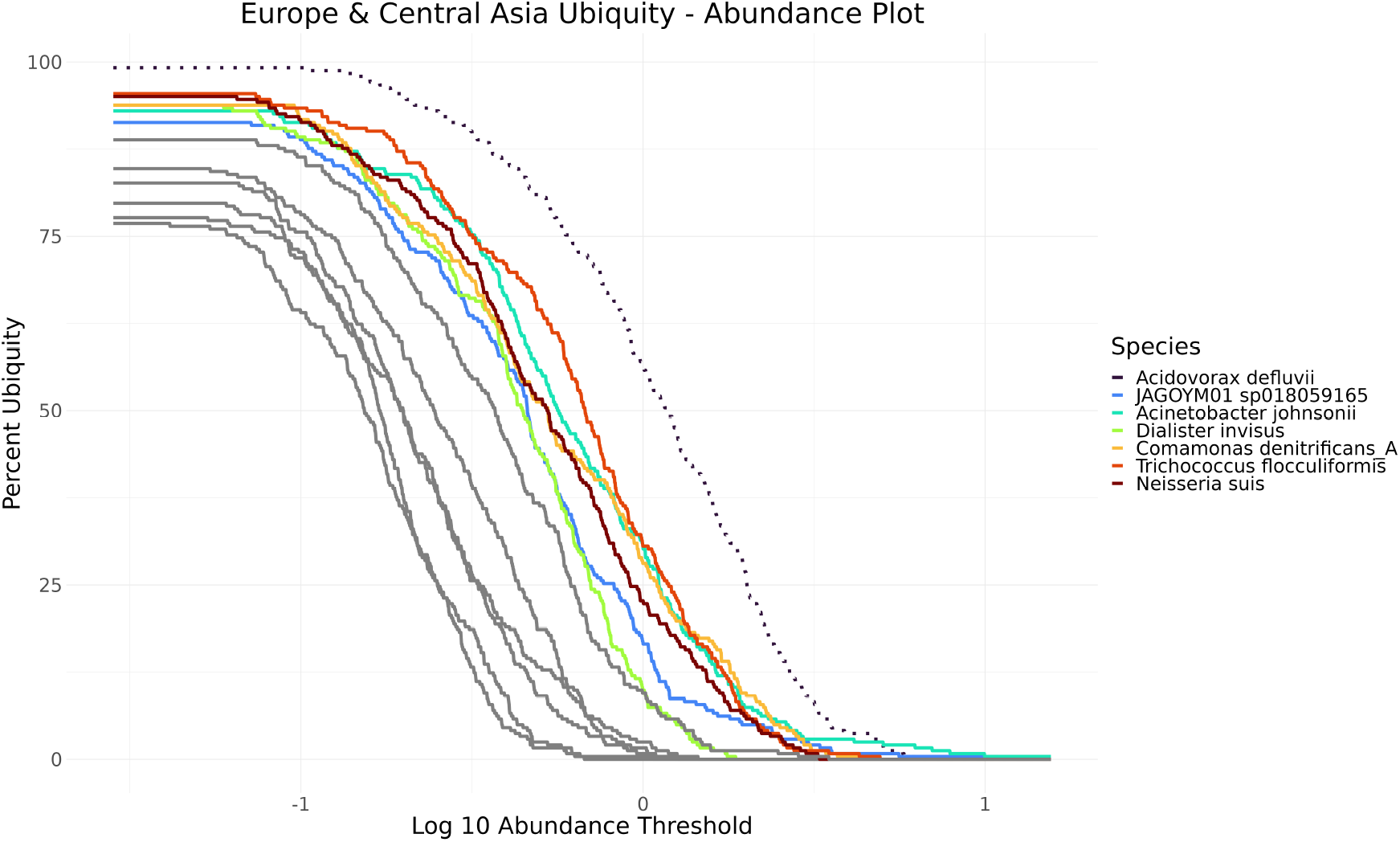
The Ubiquity - Abundance plot shows for each species its occurrence in samples (Percent Ubiquity y-axis) with a specific minimum abundance (Log10 Abundance Threshold x-axis). Only species with at least 50% ubiquity are displayed and species with more than 60% are colored.

### Machine learning guided peak memory prediction

By estimating the peak memory consumption of bioinformatics tools prior to execution, it is possible to adjust the resources requested from an infrastructure in order to minimize the needed resources per job. This approach can speed up the entire workflow by allowing more tools to run in parallel and the optimal use of the allocated cloud resources. Overall costs can be effectively utilized and even reduced by requesting only the actually necessary resources. As part of the Metagenomics-Toolkit, we use a machine learning approach (see Material and Methods for details) to improve the resource specification for two parameter sets of the MEGAHIT metagenome assembler (Figure 2, Point 3). While the Toolkit allows the assembler to be restarted on error with a user-specified higher amount of RAM, this should only be the last resort. Instead, the main goal is to predict the peak RAM usage as accurately as possible and to avoid out-of-memory crashes while using only the minimal amount of necessary RAM.

Metagenome assembly is a memory-intensive process which depends on several properties of the sequenced community. Optimization of the resource requirements of a metagenome assembler would, at best, obviate the need for dedicated rarely available high-memory hardware. Here, we train a machine learning algorithm that, for simplicity, uses the same features for two different assembler parameter settings.

We assembled 1212 metagenome datasets from environments of varying complexity, i.e. soil, biogas reactors and nasopharyngeal, twice using MEGAHIT’s default and meta-sensitive parameter settings. We selected 18 features of the datasets that had a Pearson correlation coefficient of at least 0.6 (p-value *<*0.05) with the reported peak RAM value as input for five regression-based machine learning algorithms. The best performing regressor according to our evaluation is the extremely randomized trees regressor (Supplementary Table 3).

After optimizing hyperparameters of the best estimator, we examined the feature importance reported by the regressor (Supplementary Figure 2 and 3). Based on the feature importances, we selected the number of distinct k-mers with k=21 and k=71 as features for the final model. The Nonpareil diversity index and GC content are also included in the final model but are not relevant according to feature importances and will be removed in future releases. In addition, the *Quantiles 5 k-mers* number tracks the total number of rare sequences in the dataset. The number of distinct k-mers is influenced by the diversity of the microbial community. The peak memory value depends on the complexity of the de Bruijn graph. This complexity increases with a higher number of distinct k-mers. Finally, we evaluated the best performing estimator of both models by calculating the confidence interval (95%) based on the prediction on the test dataset. The estimator of the default model has a generalization error between 3 and 5 GB RAM, while the meta-sensitive model has an error between 5 and 12 GB RAM. The maximum error reported by the confidence interval is added as a bias to the predicted value. Depending on this value, a VM flavor with the next higher memory value will be set for the execution of this dataset’s MEGAHIT assembly.

We tested the peak RAM prediction by processing 757 sewage samples. The utilized tools for the analyses can be inspected in Supplementary Table 4. As part of the assembly module, we assembled these 757 sewage samples after quality control using MEGAHIT’s meta-sensitive parameter setting. Comparing the predicted peak RAM consumption reported by Nextflow, we get a mean error of 7 GB with a standard deviation of 8 GB. In the following, we compare the amount of RAM predicted by an optimal flavor selection and four theoretical approaches (Figure 4). A scenario in which all assembler runs are started with a specific flavor is defined by the selected* modes (see different selected modes in Figure 4). In case of insufficient RAM, the assembly process runs out of memory and the flavor with the next higher RAM value is selected for the next attempt. This procedure continues until the specific sample is assembled. The optimum mode represents the amount of RAM required to assemble all samples on the first try. The theoretical approach that allocates 113 GB of RAM on the initial attempt is the one with the highest RAM consumption in total while our prediction mode results in the lowest RAM consumption compared to all naive selection approaches.

### Global occurrence of species revealed by analyzing members of the sewage core microbiome

This section presents a use case demonstrating the capabilities of the Toolkit, with a particular focus on the identification of a sewage core microbiome.

After running the per-sample workflow step, which includes assembling and binning of the 757 sewage datasets, we dereplicated all MAGs at the species level and mapped the sequence reads of each sample against the representative genomes. All results of the per-sample step are available as EMGB input files for further investigation (see Data Availability). Dereplication of all generated MAGs resulted in a set of 3473 MAGs that are at least 50% complete and at most 5% contaminated. We define the sewage core microbiome as a set of MAGs where each MAG represents a species that is present in multiple samples. Specifically, we are interested in MAGs that are present either in more than 60% of all sewage samples or in 90% of all sewage samples from a particular region and their abundance.

For all subsequent analyses, we filtered out datasets below the Q1-1.5*IQR of the Nonpareil genome coverage percentage, which resulted in the removal of five samples with low sequencing depth according to Nonpareil. We give a general overview of the organisms that meet the aforementioned core microbiome criteria by considering their occurrence in dependence of their abundance using ubiquity - abundance plots [40] (Figure 5). The first insight is that there is no MAG that could be found in all samples. This could also be due to low sequencing depth or technical limitations in the assembly or binning procedures. The MAGs present in more than 60% of all samples belong to ten species. Based on the literature (see Supplementary Table 1, column links for details), all species were found either in samples from sewage or wastewater treatment plants, with the exception of *Dialister invisus*, which was isolated from human oral cavity samples.

In Figure 5, it can be observed that the curves can cross. This occurs, for example, when a high-ubiquity, low-abundance species is compared to a low-ubiquity, high-abundance species in a subset of samples. One example is the species *Acinetobacter defluvii*, represented by a dotted line, as the species with the highest abundance in 11.3% of all samples. *Acinetobacter defluvii* is particularly common in samples taken in Europe and Central Asia (Figure 6), where it occurs in 99.1% of all samples and is the most abundant species in 29.7% of all datasets. Another continent where *Acinetobacter defluvii* occurs in many samples is North America with 83.5%. In 13.9% of all North American samples this species featured the highest abundance.

Considering the regional differences, regional core microbiomes on a continental scale were examined based solely on ubiquity. Here we applied an approach similar to the “range-through” approach described by Neu et al. [51]. We search for species that are present in more than 90% of all samples from a country and only include these species in the next step. Only if a species could be detected in 80% of all available countries in a region, we define it as part of the core microbiome of that region. Using this strategy, the following possible members of a core microbiome were detected (Table 2). *Neisseria suis* in combination with *Acidovorax defluvii* are members of the core microbiome of sewage samples in the Europe and Central Asia region and *Dialister invisus* is a core microbiome member in North America.

**Table 2.** List of organisms that are present in more than 90% of all samples from a country and their percentage ubiquity in the country grouping according to the World Bank. Only species with the highest ubiquity per grouping or with a ubiquity greater than 80% are listed.

| Country Grouping<br>according to World Bank | Species name<br>according to GTDB | Percentage of countries<br>of a World Bank grouping<br>where the organism<br>could be detected | Checkm<br>Completeness<br>(%) | Checkm<br>Contamination<br>(%) |
| --- | --- | --- | --- | --- |
| Europe & Central Asia | *Acidovorax defluvii* | 94.87 | 74.95 | 2.39 |
|  | *Neisseria suis* | 82.05 | 77.59 | 0 |
| North America | *Dialister invisus* | 100 | 65.52 | 0 |
| Sub-Saharan Africa | *Neisseria suis* | 45.45 | 77.59 | 0 |
| Middle East & North Africa | *Desulfobulbus sp017998195* | 71.42 | 98.81 | 0 |
|  | *Comamonas denitrificans_A* | 71.42 | 91.71 | 2.27 |
|  | *JAGPMH01 sp018052945* | 71.42 | 99.28 | 0.63 |
|  | *Neisseria suis* | 71.42 | 77.59 | 0 |
| East Asia & Pacific | *Escherichia coli* | 38.46 | 98.57 | 0.94 |
| Latin America & Caribbean | *Neisseria suis* | 53.84 | 77.59 | 0 |

Source: This table lists organisms present in more than 90% of samples from a country.

**Table 3.** Feature comparison between the five workflows MetaGEM, MetaWrap, MUFFIN, SqueezeMeta, and nf-core/MAG. The features listed are either Toolkit features that are implemented by at most 2 of the 5 workflows, or features that are not implemented by the Toolkit but are implemented by at least 3 of the other 5 workflows.

| Features | Implemented by the Metagenomics-Toolkit | Number of workflows that implement this feature | Toolkit features implemented by at most 2 of 5 workflows | Features not implemented by the Toolkit but by at least 3 of 5 other workflows |
| --- | --- | --- | --- | --- |
| **Sequence Data Input** |  |  |  |  |
| Assembly | ✗ | 3 |  | Missed |
| Single End Reads | ✗ | 3 |  | Missed |
| Long Read (Nanopore) | ✓ | 2 | ✓ |  |
| **Analysis Options** |  |  |  |  |
| Co-Assembly | ✗ | 3 |  | Missed |
| Dereplication | ✓ | 0 | ✓ |  |
| Plasmid Detection | ✓ | 0 | ✓ |  |
| Genome-Scale Metabolic Modelling | ✓ | 1 | ✓ |  |
| Fragment Recruitment | ✓ | 0 | ✓ |  |
| Pathway Annotation (using KEGG or similar) | ✓ | 2 | ✓ |  |
| Antibiotic Resistance | ✓ | 0 | ✓ |  |
| Contig/Gene Based Taxonomy (e.g. kraken) | ✓ | 2 | ✓ |  |
| Co-Occurrence | ✓ | 0 | ✓ |  |
| Process Unbinned Contigs | ✓ | 2 | ✓ |  |
| **Data Handling** |  |  |  |  |
| Aggregate multiple samples as a separate step | ✓ | 0 | ✓ |  |
| Optimized Database Management<sup>1</sup> | ✓ | 0 | ✓ |  |
| S3 Usage | ✓ | 2 | ✓ |  |
| Database extensibility<sup>2</sup> | ✓ | 1 | ✓ |  |
| **Other Functionality** |  |  |  |  |
| SRA Processing | ✓ | 0 | ✓ |  |
| Machine Learning Guided Parameter Estimation | ✓ | 0 | ✓ |  |
| Allow separate Workflows to call | ✓ | 2 | ✓ |  |
| HPC suited | ✗ | 5 |  | Missed |
| CI Tested | ✓ | 2 | ✓ |  |
| Visualisation Platform | ✓ | 2 | ✓ |  |

Note: This table lists features implemented by various workflows and their comparison.
<sup>1</sup>DB Download, DB integrity check, DB stored in scratch dir.
<sup>2</sup>User can provide databases.

In Latin America and the Caribbean, South Asia, Middle East and North Africa, Sub-Saharan Africa and East Asia, and Pacific region, the most ubiquitous species occurs in 53.84%, 60%, 72.42%, 45.45% and 38.46% of all countries, respectively (Table 2).

## Discussion

In this work, we presented the Metagenomics-Toolkit, a scalable workflow that provides fully reproducible results, offers different analysis methods and can process datasets on single workstations, but is particularly capable of handling large amounts of data in SLURM based clusters hosted on cloud infrastructures. The utilized Nextflow DSL2 allows code to be structured into modules, improving code readability and reusability. Hence, users can intuitively select only certain modules and steps to tailor a Toolkit run according to their preferences. The division of the workflow into a per-sample and an aggregation step allows the distribution of the first part to multiple cloud sites to complete the analysis in an acceptable time frame, especially in demanding cases with thousands of samples. The optimized input and database handling has significantly improved the speed and reliability of the workflow, while reducing the risk of bottlenecks in the file system. Computed results are more consistent, reproducible and predictable regarding disk allocation, making it easier to carry out large scale bioinformatics analyses in a cloud computing environment. However, the overall performance can be further improved by relying less on a shared file system and more on object storage. It must be further investigated which solution Nextflow offers to move the working directory to an S3 compatible object storage. To reduce the likelihood of programming errors being introduced, the Metagenomics-Toolkit is tested regularly against toy datasets and real world datasets by using Github Actions. These tests are executed for each module and a range of different combinations of modules.

We have demonstrated the usefulness of assembly peak RAM prediction when processing hundreds of datasets. It should be noted that in addition to the benefit of our prediction mode enabling the lowest RAM consumption in comparison to the naive approaches, it automatically selects the VM flavor with the next higher RAM value in case of an error. Combined with error handling, the prediction mode eliminates the need for manual intervention and thereby automates the failover processing of a large number of datasets. Although in our theoretical modes, the selected medium flavor mode is close to our prediction mode, in practice, the user would have to estimate the actual RAM consumption based on experience. The error in selecting the wrong flavor based on an inaccurate predicted peak RAM consumption depends on the user defined flavor set. Setting wider RAM flavor boundaries leads to more correct flavor selections. An additional insight is that according to the feature importances (Supplementary Figure 2 and 3) of both models, the size of the dataset (number of bases) is not as important as the k-mer statistics for a correct estimation. It is to be noted that due to the bias that is added to the predicted value, also low-complexity datasets that may only require a few GB of RAM will always end up with additional GB of RAM according to the bias. This may make it impossible to run the assembly on a user workstation, i.e., a laptop. In these cases, the prediction mode can be disabled in the configuration file. While the created machine learning model only applies to MEGAHIT, the machine learning method we used to create the model allows it to be applied to other assemblers as well, such as metaSPAdes. In addition, it needs to be investigated whether the model can be further simplified without loss of accuracy.

Regarding the use case being performed, to our knowledge, it is the first analysis of the sewage core microbiome with such a large number of datasets. Regarding the core microbiome definition, it was pointed out by Neu et al. [51], that there is no single ubiquity or abundance threshold that has been used in previous studies to define a core microbiome. Therefore, we utilized ubiquity-abundance plots to give the reader an overview of a range of possible thresholds before we define our own ubiquity threshold. However, we only used ubiquity thresholds, since low abundance genomes could also be important members of a community and we don’t have any reason to exclude them when defining the core microbiome. Our analysis highlights the distribution of species worldwide and in specific regions. We examined not only the presence but also the abundance of all organisms, allowing future work to analyze the potential spread of AMR and associated microorganisms such as *E. coli*.

In general, we expect that members of the core microbiome are widespread organisms that have characteristics such as versatility in terms of nutrient uptake, stress tolerance or adaptation of defense mechanisms, and to be able to withstand seasonal effects, as samples were taken at different times of the year. One of the organisms we detected is *Acinetobacter johnsonii* which according to Jia et. al. [26] shows exceptional adaptability to occupy different environments. However, the actual characteristics of the organisms should be investigated in follow-up analyses. It should also be noted that although we present a geographical distribution of the organisms, it needs to be investigated whether the occurrence of the organisms depends on geography or on other factors.

We have shown that the Metagenomics-Toolkit is ideal for large scale and efficient analysis on cluster-based cloud systems enabling the investigation of the sewage core microbiome by processing 757 metagenome samples. The uniqueness of its feature set is discussed further in the following chapter.

### Feature comparison of existing metagenomics workflows

The Metagenomics-Toolkit offers different tools, functionalities and analyses combined with different sequence data input types. While some capabilities are already available in existing workflows, we want to highlight the novel ones in this chapter. We compared the Metagenomics-Toolkit to five metagenomics workflows, namely MetaGEM, MetaWrap, MUFFIN, SqueezeMeta, and nf-core/MAG, that meet the requirements that the workflow must be fully publicly available, represent a workflow where all parts can be executed in a single call, and be in common use at the time of writing. We compared all workflows in terms of their implemented features in four categories: “Sequence Data Input”, “Analysis Options”, “Data Handling” and “Other Features”. We only refer to two types of features: One type are features that are implemented by at most 2 out of 5 workflows (“Novel Functionality”) and the other type represents features implemented by at least 3 out of 5 workflows but not by the Toolkit (“Missing Features”) (Table 3). A comparison table with all features can be found in Supplementary Table 2.

Novel features available in the Metagenomics-Toolkit are, as described in the previous main section, the co-occurrence, fragment recruitment, plasmid detection and peak RAM consumption prediction. In contrast to most other workflows, the Metagenomics-Toolkit allows the input of datasets obtained by the Oxford Nanopore sequencing technology. However, it does not calculate hybrid assemblies by accepting short and long reads. The Metagenomics-Toolkit offers many types of analysis methods such as a search for genes predicted to mediate antibiotic resistance and genome-scale metabolic modeling of the reconstructed MAGs. In addition, the Toolkit is optimized to work directly with SRA data and to scale on cloud-based clusters. Finally, a distinguishing feature is the ability to explore metagenomic data via an interactive website with the EMGB.

Several other workflows offer co-assembly functionality, which is particularly useful for recovering MAGs with low sequence coverage. However, there is a trade-off between our implemented separate assembly combined with a dereplication approach and co-assembly. Co-assembly is expected to generate MAGs with higher completeness, and to yield more low coverage genomes. However, it results in higher contamination compared to single assemblies and subsequent dereplication [85]. It remains to be evaluated in which cases co-assembly should be preferred over our approach or in which cases a combination of both strategies could be used.

In addition to the co-assembly functionality, future enhancements will focus on the improvement of specific modules and on additional input data types such as assembled contigs, transcriptome data and a combination of short and long reads for hybrid assemblies. The co-occurrence module can be better integrated with genome-scale metabolic modeling. One example would be that detected sub-communities in the network could be better investigated according to their MRO and MIP values. In general, new modules can be introduced that provide a pangenome analysis or the analysis of viral genomes.

Finally, due to the extensive use of Docker, the Metagenomics-Toolkit needs to be adapted for use in HPC environments. Nextflow allows to easily specify other HPC-friendly container engines like Singularity [34] or Podman [23].

## Data Availability

### Code

All tools and containers that were used can be viewed by accessing the tag “0.3.0-rc.15” of the https://github.com/metagenomics/metagenomics-tkrepository.

### Configuration

The actual configuration with a list of used databases can be found in this repository: https://github.com/metagenomics/wastewater-study

### Sewage Analysis

The Toolkit output of all sewage datasets is publicly available via the S3 link s3://wastewater using the endpoint url https://openstack.cebitec.uni-bielefeld.de:8080.

### EMGB

EMGB inputs of every dataset can be downloaded via the S3 link s3://wastewater-emgb using the endpoint url https://openstack.cebitec.uni-bielefeld.de:8080. Further details regarding sewage datasets in EMGB can be found in the wastewater-study repository: https://github.com/metagenomics/wastewater-study

