## Supplemental Figures for "Metagenomics-Toolkit: The Flexible and Efficient Cloud-Based Metagenomics Workflow featuring Machine Learning-Enabled Resource Allocation"

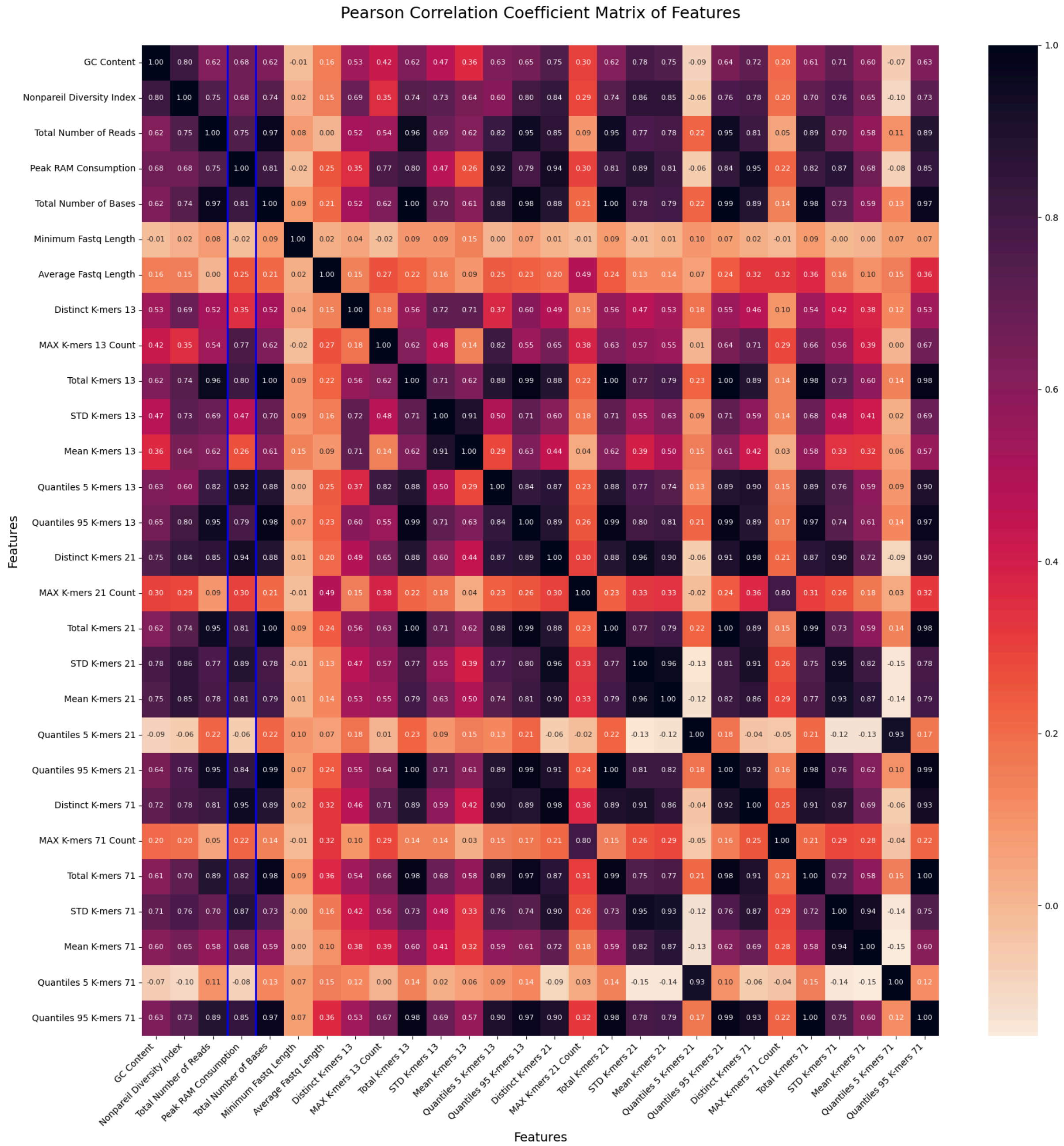


Figure 1: Pearson correlation matrix of possible features for a machine learning algorithm.

Explanation

- MAX K-mer * Count: The maximum of the total number of k-mers per k-mer frequency.
- Total K-mers *: The total number of k-mers.
- STD K-mers *: Standard deviation of total number of k-mers per k-mer frequency.
- Mean K-mers *: Mean of the total number of k-mers per k-mer frequency.
- Distinct K-mers *: Number of distinct k-mers.
- Quantiles 5 K-mers *: The sum of the lowest 5% of the total number of k-mers per k-mer frequency.

- Quantiles 95 K-mers *: The sum of the highest 5% of the total number of k-mers per k-mer frequency.

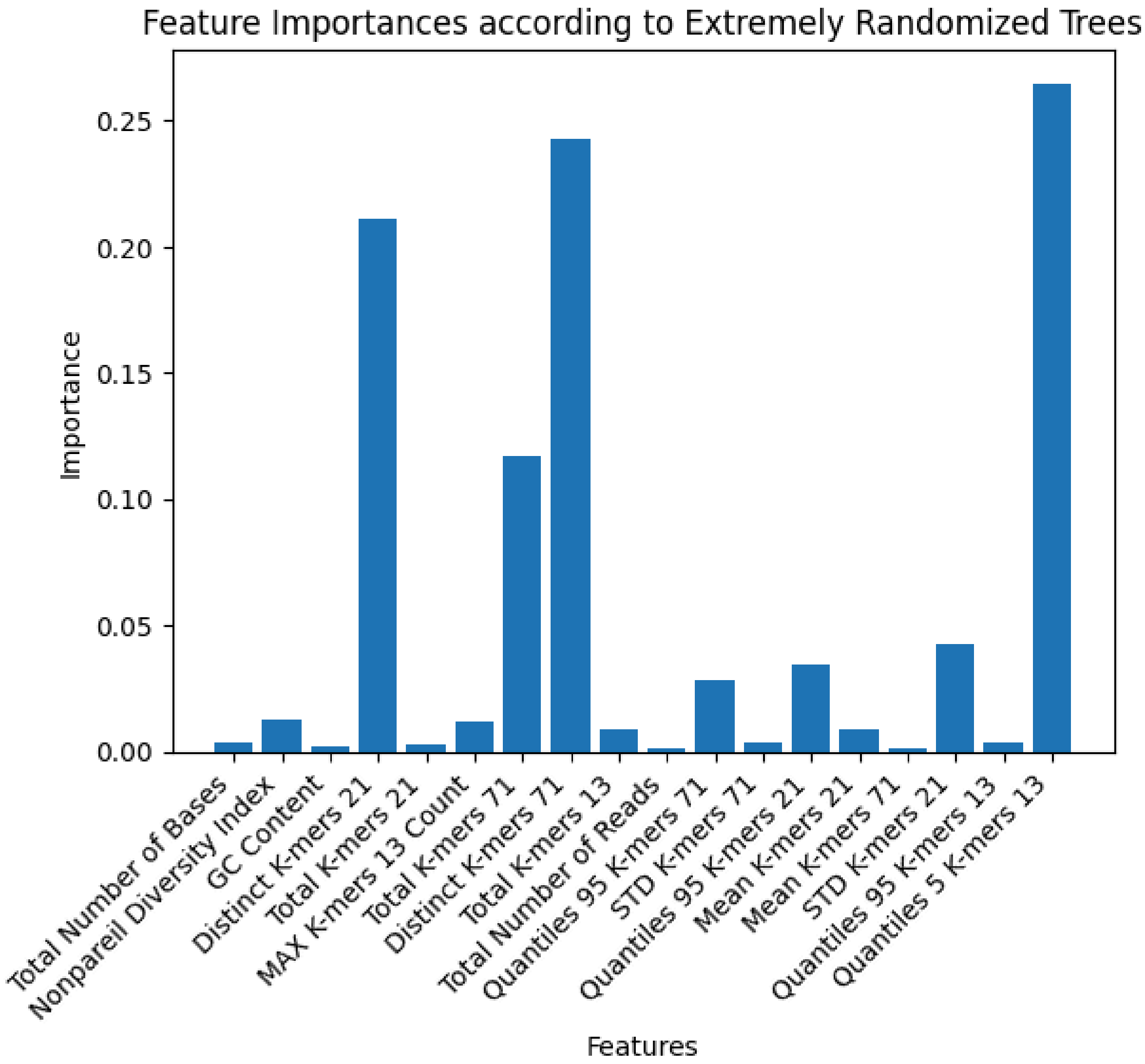


Figure 2: Feature importances according to Extremely Randomized Tree approach based on Megahits default parameters.

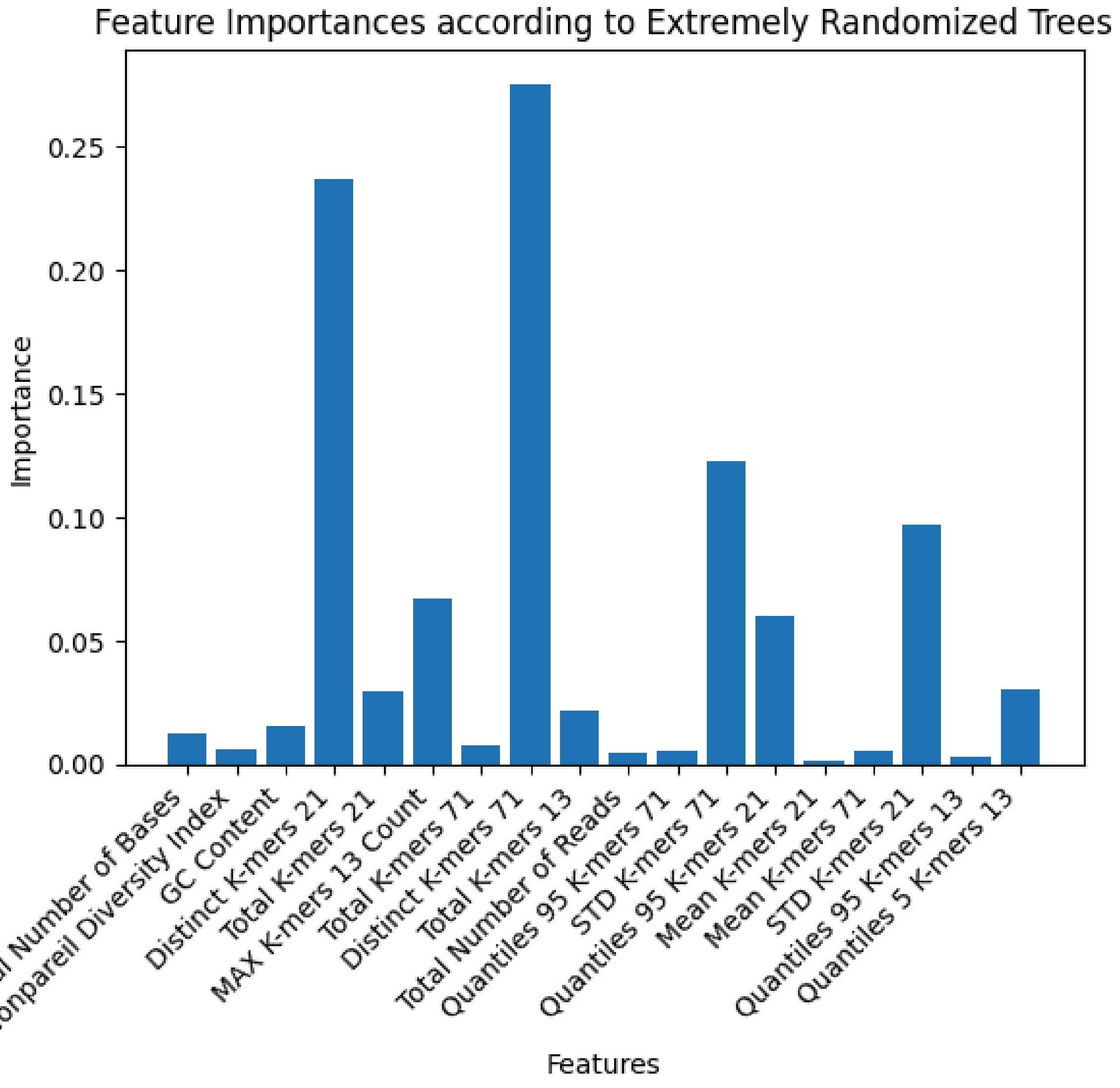


Figure 3: Feature importances according to Extremely Randomized Tree approach based on Megahits meta-sensitive parameters.

```
AGGREGATED
└── 1
    └── fragmentRecruitment
        └── 0.1.0
            └── mash
                └── paste

ERR4678561
└── 1
    ├── annotation
    │   └── 0.3.0
    │       ├── mmseqs2
    │       │   ├── bacmet20_experimental
    │       │   ├── bacmet20_predicted
    │       │   ├── kegg
    │       │   ├── ncbi_nr
    │       │   └── vfdb
    │       ├── mmseqs2_taxonomy
    │       │   └── gtdb
    │       ├── prokka
    │       │   └── logs
    │       └── rgi
    │           └── logs
    ├── assembly
    │   └── 1.2.1
    │       ├── megahit
    │       └── predictFlavor
    ├── binning
    │   └── 0.5.0
    │       ├── contigCoverage
    │       ├── contigMapping
    │       ├── genomeCoverage
    │       │   └── logs
    │       ├── metabat
    │       └── readMappingQuality
    ├── config
    │   └── 0.1.0
    │       └── parameters
    ├── fragmentRecruitment
    │   └── 0.1.0
    │       └── mashScreen
    ├── magAttributes
    │   └── 1.0.1
    │       ├── checkm
    │       │   └── logs
    │       └── gtdb
    │           └── logs
    ├── metabolomics
    │   └── 0.1.0
    │       ├── carveme
    │       │   └── logs
    │       ├── gsmmJson
    │       │   └── logs
    │       ├── gsmmTsv
    │       │   └── logs
    │       └── memote
    │           └── logs
    ├── plasmids
    │   └── 1.0.0
    │       ├── MobTyper
    │       │   └── logs
    │       ├── PLSDB
    │       │   └── logs
    │       ├── PlasClass
    │       │   └── logs
    │       ├── SCAPP
    │       │   ├── coverage
    │       │   ├── logs
    │       │   └── readMapping
    │       ├── ViralVerifyPlasmid
    │       │   └── logs
    │       └── filtered
    └── qc
        └── 0.3.0
            ├── fastp
            ├── jellyfish
            └── nonpareil
```

Figure 4: Example directory output structure of one Toolkit sample run with the aggregation step.

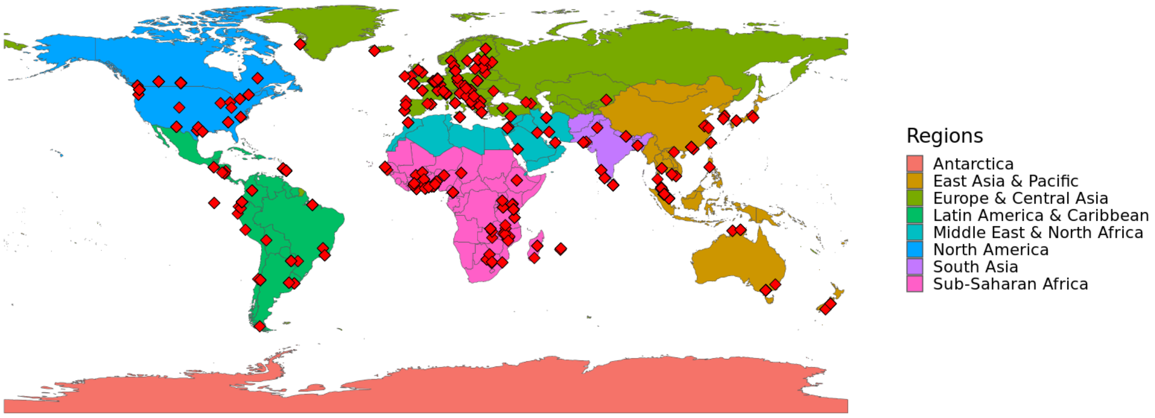


Figure 5: World map colored according to the World Bank.

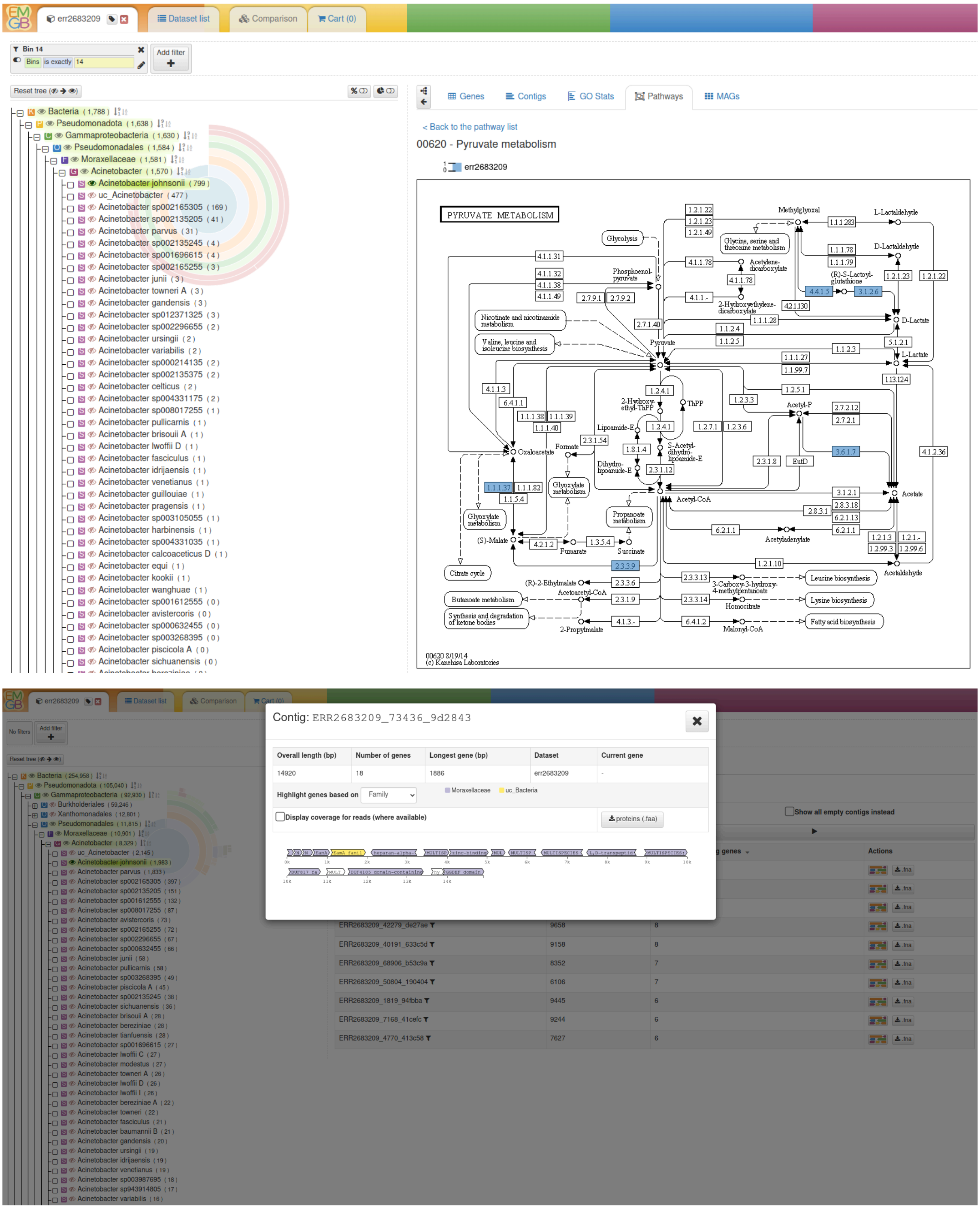

Figure 6: Screenshots of the Exploratory MetaGenome Browser (EMGB).
*The upper screenshot displays a filtered set of genes and their occurrence in the Pyruvate Metabolism. The screenshot at the bottom shows genes that can be inspected by the contig viewer.*
